# Initiator tRNA Genes Template the 3’CCA End at High Frequencies in Bacteria

**DOI:** 10.1101/035220

**Authors:** David H. Ardell, Ya-Ming Hou

## Abstract

While the CCA sequence at the mature 3’ end of tRNAs is conserved and critical for translational function, a genetic template for this sequence is not always contained in tRNA genes. In eukaryotes and archaea, the CCA ends of tRNAs are synthesized post-transcriptionally by CCA-adding enzymes. In bacteria, tRNA genes template CCA sporadically. In order to understand variation in how prokaryotic tRNA genes template CCA, we re-annotated tRNA genes in the tRNAdb-CE database. Among 132,129 prokaryotic tRNA genes, initiator tRNA genes template CCA at the highest average frequency (74.1%) over all functional classes except selenocysteine and pyrrolysine tRNA genes (88.1% and 100% respectively). Across bacterial phyla and a wide range of genome sizes, many lineages exist in which predominantly initiator tRNA genes template CCA. Preferential retention of CCA in initiator tRNA genes evolved multiple times during reductive genome evolution in Bacteria. Also, in a majority of cyanobacterial and actinobacterial genera, predominantly initiator tRNA genes template CCA. We suggest that cotranscriptional synthesis of initiator tRNA CCA 3’ ends can complement inefficient processing of initiator tRNA precursors, “bootstrap” rapid initiation of protein synthesis from a non-growing state, or contribute to an increase in cellular growth rates by reducing overheads of mass and energy to maintain nonfunctional tRNA precursors. More generally, CCA templating in structurally non-conforming tRNA genes can afford cells robustness and greater plasticity to respond rapidly to environmental changes and stimuli.

## INTRODUCTION

All active tRNA molecules must contain a CCA sequence at the 3’-end as the site for amino acid attachment and for interaction with the ribosome during protein synthesis (Betat, Rammelt et al. 2010, Vortler and Morl 2010, Betat and Morl 2015). While essential for tRNA activities, the CCA sequence is generally not encoded in tRNA genes but is added post-transcriptionally. Exceptions are found in bacteria, where some tRNA genes contain a template of the CCA sequence for direct synthesis at the time of transcription. However, CCA-templating is not necessarily conserved among tRNA genes with different functional identities or among bacterial species across different phyla. To explore whether there is potential selective pressure for tRNA genes to template CCA in bacteria, we undertook a reannotation of publicly available tRNA gene data.

One source of error in the annotation of tRNA genes concerns the functional classification of genes for tRNAs with CAU anticodons. These include genes for both the initiator and elongator tRNA^Met^ and specific elongator tRNA^Ile^_CAU_ isoacceptors in bacteria and archaea. In the latter case, transcribed CAU anticodons are post-transcriptionally modified to distinguish them from the unmodified CAU anticodons of cytosolic tRNA^Met^ (Suzuki and Miyauchi 2010). However, currently available tRNA gene-finders annotate all three classes as elongator tRNA^Met^ genes (Ardell 2010). The TFAM tRNA functional classifier, which uses profile-based models of whole tRNA sequences (Ardell and Andersson 2006, Tåquist, Cui et al. 2007), can differentiate all three tRNA functional classes with generally high specificity and sensitivity (Silva, Belda et al. 2006). However, the tRNA^Ile^_CAU_ class evolves more rapidly than other classes, so that even though the TFAM 1.4 Proteobacterial-specific model generalizes well to some other Bacterial phyla, this model does not generalize well to all (Freyhult, Cui et al. 2007). An alternative TFAM model (Amrine, Swingley et al. 2014), for just genes for tRNAs with CAU anticodons, is based on a custom annotation of such genes in a wide sampling of bacterial taxa (Silva, Belda et al. 2006). Although this alternative model is imperfect in its sensitivity and specificity (Silva, Belda et al. 2006), as discussed further below, its performance is satisfactory and suitable for the present study.

Here we apply the alternative “Silva TFAM model” to improve the functional annotation of tRNA genes with CAU anticodons in the high quality public database tRNAdb-CE (Abe, Ikemura et al. 2009, Abe, Inokuchi et al. 2014). In our analysis, we found that genes for the initiator class of tRNAs across the bacterial domain consistenly template CCA with significantly higher frequencies than elongator tRNA genes. This CCA-templating can provide unique advantages to initiator tRNA for rapid maturation, aminoacylation, and initiation of protein synthesis.

## RESULTS

### Functional Reannotation of Bacterial Genes in tRNAdb-CE v.8

The tRNAdb-CE v0.8 database uses TFAM 1.4 for functional classification of bacterial tRNA genes (as described in http://trna.ie.niigata-u.ac.jp/trnadb/method.html). However, the Proteobacterial model for the tRNA^Ile^_CAU_ elongator class that comes with TFAM 1.4 does not generalize well to all bacterial phyla (Freyhult, Cui et al. 2007). Therefore, we reannotated 9,914 bacterial genes for tRNAs with CAU anticodons in tRNAdb-CE v0.8 using the more general Silva TFAM model derived from the analysis. This model is also provided as supplementary data in the present work. By applying the Silva TFAM model, we revised the functional classification of 4,362 of 9,914 genes (≈ 43.9%). Reclassification frequencies are presented in Table 1, showing that most of the changes involve reclassification of genes from tRNA^Met^ to initiator tRNA^fMet^ or to tRNA^Ile^_CAU_. Reannotated data are provided in supplementary materials.

**Table 1:**
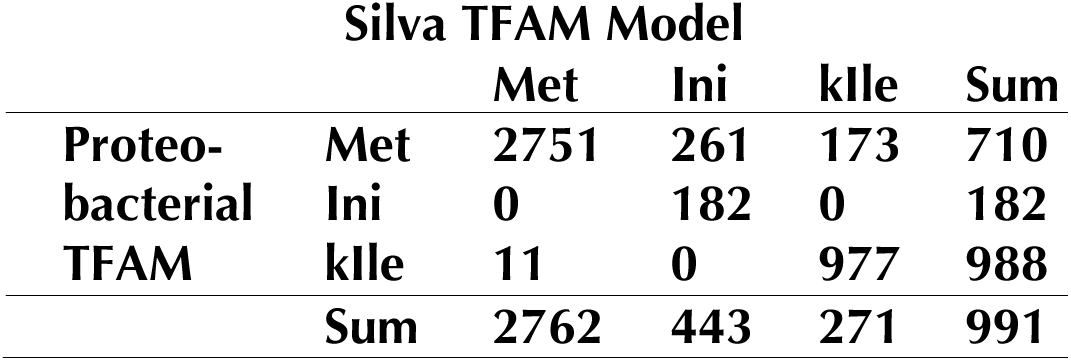
Reannotation of bacterial tRNAs with CAU anticodons in tRNAdb-CE v.8 using a custom model based on the analysis of (Silva et al., 2006)

### Structural Reannotation of Bacterial Genes in tRNAdb-CE v.8

A well-designed feature of the tRNAdb-CE v0.8 data model lies in that its gene records contain not only annotated gene sequences but also ten bases of genomic context both up and downstream. Inspection of tRNAdb-CE v0.8 data revealed multiple genes with an annotated 3’-end sequence other than CCA, followed by 3’-trailer sequences that begin with the sequence CCA. To confidently assess whether these genes might template a 3-CCA-end for their gene products, we assigned Sprinzl coordinates (Sprinzl, Horn et al. 1998) to these bases for each gene sequence. These coordinates were not provided in tRNAdb-CE v0.8. We did this by implementing a dynamic programming algorithm to optimize base-pairing of the acceptor 3’-end region against the database-annotated 5’-end. Although our acceptor-end annotations were almost always identical with those annotated in the database, they enabled us to confidently and consistently assign Sprinzl coordinates to the 3’-end region of each gene. Using this technique, we annotated an additional 2,866 bacterial tRNA genes out of 129,989 (or 2.2%) records as containing the CCA template at the 3’-end in the sequence framework of Sprinzl coordinates 74–76.

To clarify why we could identify an additional 2,866 tRNA genes in tRNAdb-CE v0.8 that template CCA, we ran tRNA gene-finding programs on the database records. We used ARAGORN v1.0 (Laslett and Canback 2004) and tRNAscan-SE v.1.23 (Lowe and Eddy 1997) in default eukaryotic tRNA gene-finding mode, and tRNAscan-SE v.1.23 in Bacterial mode (with the -B option). We found that tRNAscan-SE v.1.23, when run in its default eukaryotic gene-finding mode, never annotates nucleotides at positions 74–76 irrespective of sequence.

An exception to this rule was with selenocysteine tRNA genes, for which tRNAscan-SE in eukaryotic mode does annotate positions 74–76 if they contain the CCA sequence. From this observation we conclude that a likely cause of misannotations in tRNAdb-CE is user error in genome annotation pipelines. This is particularly notable when users of tRNAscan-SE use its default eukaryotic gene-finding mode on prokaryotic genomes. Such errors may then be incessently propagated in public and private databases.

### Frequencies of CCA-templating in Bacterial tRNA Genes

With our reannotated tRNAdb-CE data in hand, we calculated frequencies of CCA-templating in tRNA genes across different tRNA functional classes and taxonomic groupings as defined by NCBI Taxonomy. Figure 1 visualizes our data summarized by prokaryotic genus. Prokaryotic clades exhibit all four possible patterns: 1) all tRNAs genes template CCA, 2) few or no tRNA genes template CCA, 3) primarily initiator genes template CCA, or — most rarely — 4) primarily elongator tRNA genes template CCA.

**Figure 1.**
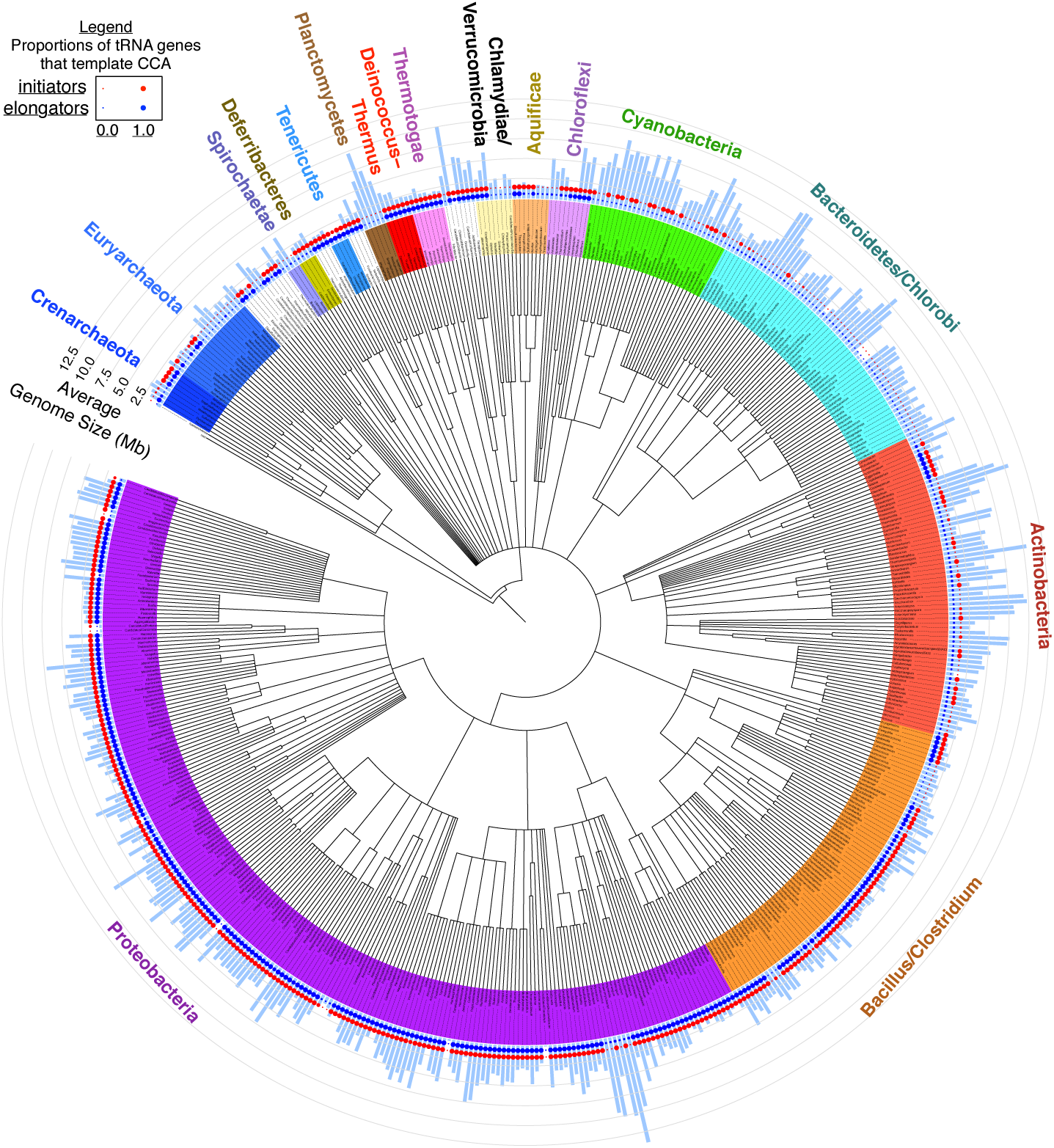
Summarized frequencies at which elongator tRNA genes and initiator tRNA genes template 3’-CCA against average genome size in different prokaryotic genera. NCBI-Taxonomy based cladogram of prokaryotic genera in tRNAdb-CE v.8 showing average genome size (radial light blue bars) and average fractions at which elongator tRNA genes template 3’-CCA (blue circles) and initiator tRNA genes template 3’-CCA (red circles).

The five best-sampled phyla in our dataset, as defined by number of distinct genera with at least one genome sequenced, are Proteobacteria, Bacillus/Clostridium, Actinobacteria, Bacteroidetes/Clorobi, and Cyanobacteria. These five phyla exhibit three of four patterns described above in a strikingly consistent pattern by phylum. Practically all tRNA genes template CCA in Proteobacteria and Bacillus/Clostridium, except in certain reduced genomes, most of which template CCA only in initiator tRNA genes, or in no tRNA genes at all. In Cyanobacteria and Actinobacteria, on the other hand, primarily only the initiator tRNA genes template CCA, with certain exceptions. For example, a clade of Actinobacteria with relatively small genomes exists in which both initiator and elongator tRNA genes template CCA at high frequencies. In the Bacteroidetes/Chlorobi group, most tRNA genes do not template CCA, except for one lineage, the Solitalea, in which only initiator tRNA genes template CCA. In all five of the most-sampled phyla, there exist both small and moderately-sized genomes in which only initiator tRNA genes template CCA, or no tRNA genes at all template CCA. Certain Myxococcales, among the Deltaproteobacteria, are exceptional in having among the largest genomes that we observed and yet no tRNA genes or only initiator tRNA genes template CCA.

Less-sampled phyla are also quite heterogeneous in our dataset. In the Thermotogae, Deinococcus/Thermus and Tenericutes, all tRNA genes template CCA. Spirochaetae do not template CCA in any genes, while in Deferribacteres, only initiator tRNA genes template CCA. In the most rare pattern we observed, in only a few archaeal or bacterial genera, primarily elongator tRNA genes and not initiator tRNA genes template CCA.

In order to better visualize these data down to individual genomes and separating different elongator classes, we created an interactive javascript-based taxonomic navigator for our results visualized with heatmaps in any ordinary web browser. The full interactive data navigator is available as supplementary materials to this work. A static view on these data is also provided as a searchable PDF in supplementary materials. Figure 2 presents a snapshot from this browser with some notable detailed results for Bacterial tRNA genes. The figure shows columns of frequency data, corresponding to functional classes of tRNA genes, sorted left to right by decreasing average frequency at which tRNA genes template CCA over all prokaryotic genomic sequences in our sample. This analysis reveals that initiator tRNA genes (labeled as Ini) template CCA at the highest frequency (74.1%), versus 66.2% for elongator tRNA genes generally in prokaryotes. Bacterial initiator tRNA genes template CCA at a frequency of 74.7%, second only to selenocysteinyl tRNA genes (tRNA^Sec^, Sec = selenocysteine), with a frequency of 89.2%. We also found that genes for tRNA^Asp^ and tRNA^Asn^ template CCA at the highest frequencies among all canonical elongator tRNA genes. Below we describe some of the notable results shown in Figure 2.

**Figure 2.**
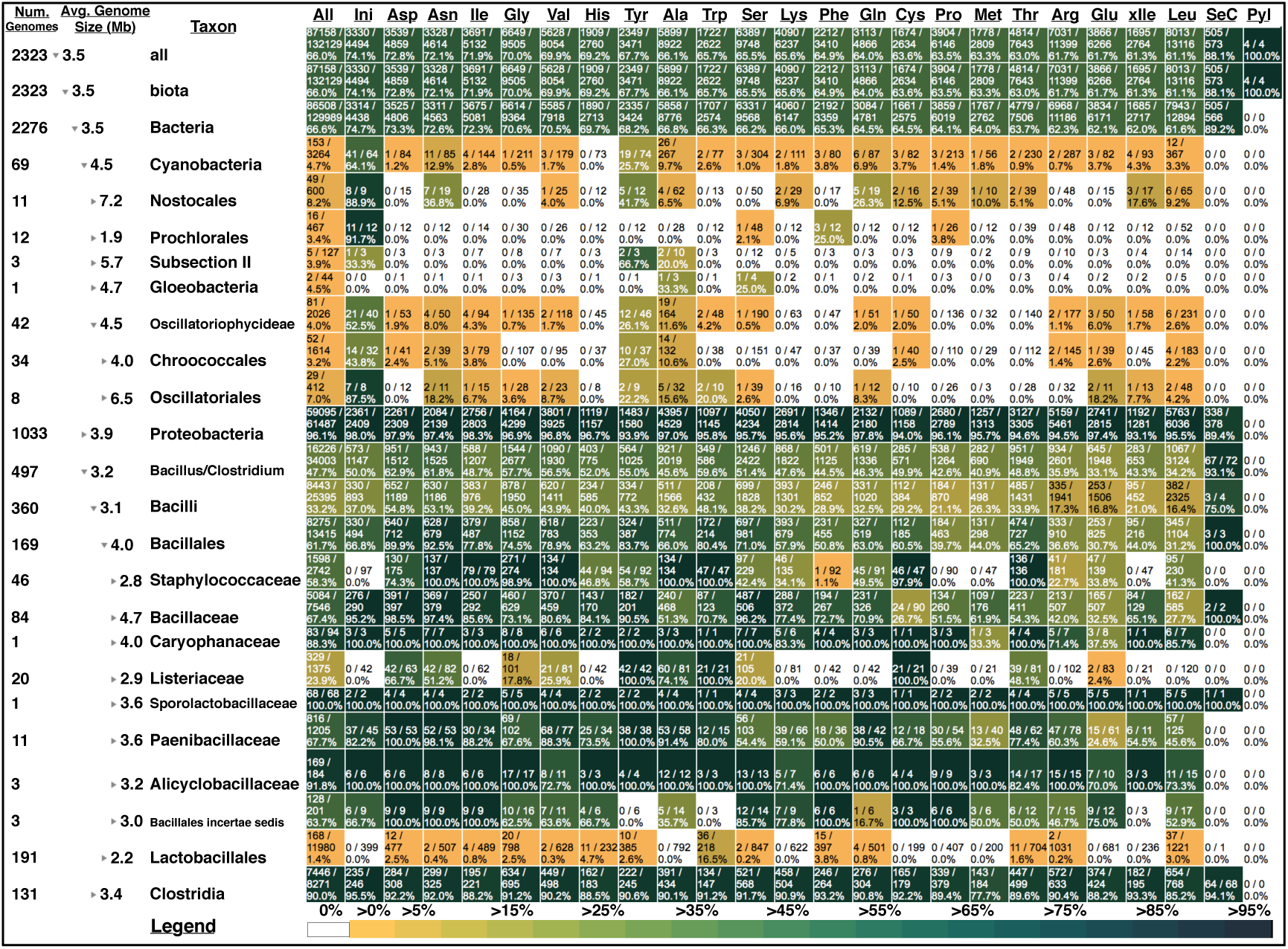
Summarized genome size and CCA frequency data in Bacterial clades broken out by tRNA functional class. Except for columns labelled “All,” “SeC,” and “Pyl,” all columns of frequency data are sorted in decreasing order from left to right in frequency at which tRNA genes template CCA over all prokaryotic genomes that we sampled. Clades are defined as in NCBI Taxonomy. Column labels correspond to IUPAC three-letter amino acid charging identity except for “Ini” (initiators) and “xIle” (AUA-codon-reading isoleucine isoacceptors). “All” summarizes frequency data over all tRNA classes.

***Cyanobacteria***. Among bacterial phyla we observed, Cyanobacteria have the most striking and consistent pattern in which specifically initiator and not elongator tRNA genes template CCA. The overall rates are 64.1% for initiator tRNA genes versus 25.7% for the next highest gene class, which are elongator tRNA^Tyr^ genes. But different cyanobacterial lineages exhibit considerable variation in this trait. For example, among Prochlorales and Nostocales genomes — comprising both the smallest and largest average genome sizes, respectively — the frequencies at which initiator tRNA genes template CCA are 91.7% and 88.9%, while elongator tRNA genes template CCA at only 3.4% and 8.2% respectively. In 11 out of 12 of *Prochlorococcus* genomes and 9 out of 13 *Synechococcus* genomes, only initiator tRNA genes template CCA. Initiator tRNA genes template CCA at very different rates in sister orders Oscillatoriales and Chroococcales within subclass Oscillatoriophycideae: 87.5 and 43.8% respectively.

The Cyanobacteria are also unusual in that different strains and groups feature specific elongator tRNA gene classes that also template CCA at intermediate rates (above 10%) while other elongator classes template CCA at lower rates (below 10%). Usually, if in any one genome the initiator tRNA gene or genes template CCA, at least one elongator tRNA gene class will also template CCA at an intermediate rate. The elongator gene class that templates CCA most consistently across the phylum is the tRNA^Tyr^ gene class. In *Nostocales*, tRNA^Tyr^ genes template CCA at a frequency of (41.7%), while tRNA^Asn^ and tRNA^Gln^ elongator genes also template CCA at a high relative rate (36.8% and 23.7%).

*Proteobacteria*. All proteobacterial tRNA genes generally template CCA at consistently high rates: 96.1% overall (Figure 2). Yet proteobacterial initiator tRNA genes template CCA at 98.0%, significantly higher than proteobacterial elongators (*χ*2 = 23.625, d.f. = 1, *p <* 10^−4^ by Fisher’s Exact Test with a Yates correction). Closer examination of the proteobacterial variation (supplementary materials) reveals that while many free-living proteobacteria template CCA at high rates, endosymbiotic γ-proteobacteria and α-proteobacteria with reduced genomes show similar patterns to those described above for cyanobacteria with reduced genomes. In these cases, initiator tRNA genes appear to be the only class to consistently template CCA, while several elongator classes also template CCA. For example, in most *Buchnera aphidicola* genomes, about eight or nine additional elongator tRNA classes template CCA at intermediate to high rates while other classes do not template CCA, as previously reported (Hansen and Moran 2012). However, not previously reported is that in all *Buchnera* strain genomes except one, initiator tRNA genes always template CCA. Furthermore, like in the Cyanobacteria, in the smallest of the *Buchnera* genomes, only initiator tRNA genes template CCA. This same pattern holds in other endosymbiotic γ-proteobacteria genomes such as Ca. *Blochmannia, Wigglesworthia, Glossina*, Ca. *Baumannia*, Ca. *Carsonella*, Ca. *Portiera*, as well as α-proteobacteria endosymbionts such as *Wolbachia*. In contrast, among the smallest γ-proteobacterial genomes like Ca. *Hodgkinia*, none of the tRNA genes template CCA.

*Other bacterial phyla*. Many diverse genera and classes of bacteria preferentially template CCA in their initiator tRNA genes (Supplementary files). Examples include *Geobacillus, Thermoaerobacter, Ruminococcus, Thermomicrobiales, Deferribacter*, Thermodesulfobacteria, *Mycobacterium, Propionibacterium, Frankia*, and *Bifidobacterium*. As shown in Figure 2, within the Bacillus/Clostridium phylum, frequency variation in this genomic trait also extensive. An unusual pattern is found in the pathogenic Staphylococcaeceae and Listeraceae families, and also the Lactobacillales, which contain both pathogens and non-pathogens, in which initiator tRNA genes never template CCA, even while elongator tRNA genes do template CCA at intermediate rates. For example, in Staphylococcaceae about 60% of elongator tRNA genes template CCA and in Listeraceae about 24% of elongator tRNA genes template CCA, while in Lactobacillales, 1.4% of elongator genes template CCA. Yet among the 257 genome representatives of these three families in our dataset, not one initiator tRNA gene templates CCA.

*Archaea*. We found no need for structural or functional reannotation of archaeal tRNA genes in tRNAdb-CE v.8. Figure 3 presents a snapshot from this browser with some of our most notable results for archaeal tRNA genes. While there are fairly high frequencies of CCA-templating in archaeal tRNA genes overall, at 30.4%, we found that initiator tRNA genes in Archaea do not template CCA at any especially high frequency among tRNA genes, which presents a major difference from bacteria. Other than this, we observed extensive phyletic variation in this trait across Archaea. Crenarchaeota tRNA genes template CCA at a rate of 50.5%, while Euryarchaeota tRNA genes template CCA at about half of that rate. Within Crenarchaeota, tRNA genes in the Sulfolobales template CCA at 3.8%, but in the Desulforococcales this rate is 84.8%. All four tRNA^Pyl^ (Pyl = pyrrolysine) genes template CCA in the Methanomicrobia. Contrary to the generalization that Archaea and Eukarya do not template CCA, there exist lineages in both the Crenarchaeota and Euryarchaeota in which all or nearly all tRNA genes template CCA, for example in the Desulfurococcales, Protoarchaea, and Methanopyri. Although variation exists across tRNA functional classes in a phyletic pattern, no obvious overall pattern emerges.

**Figure 3.**
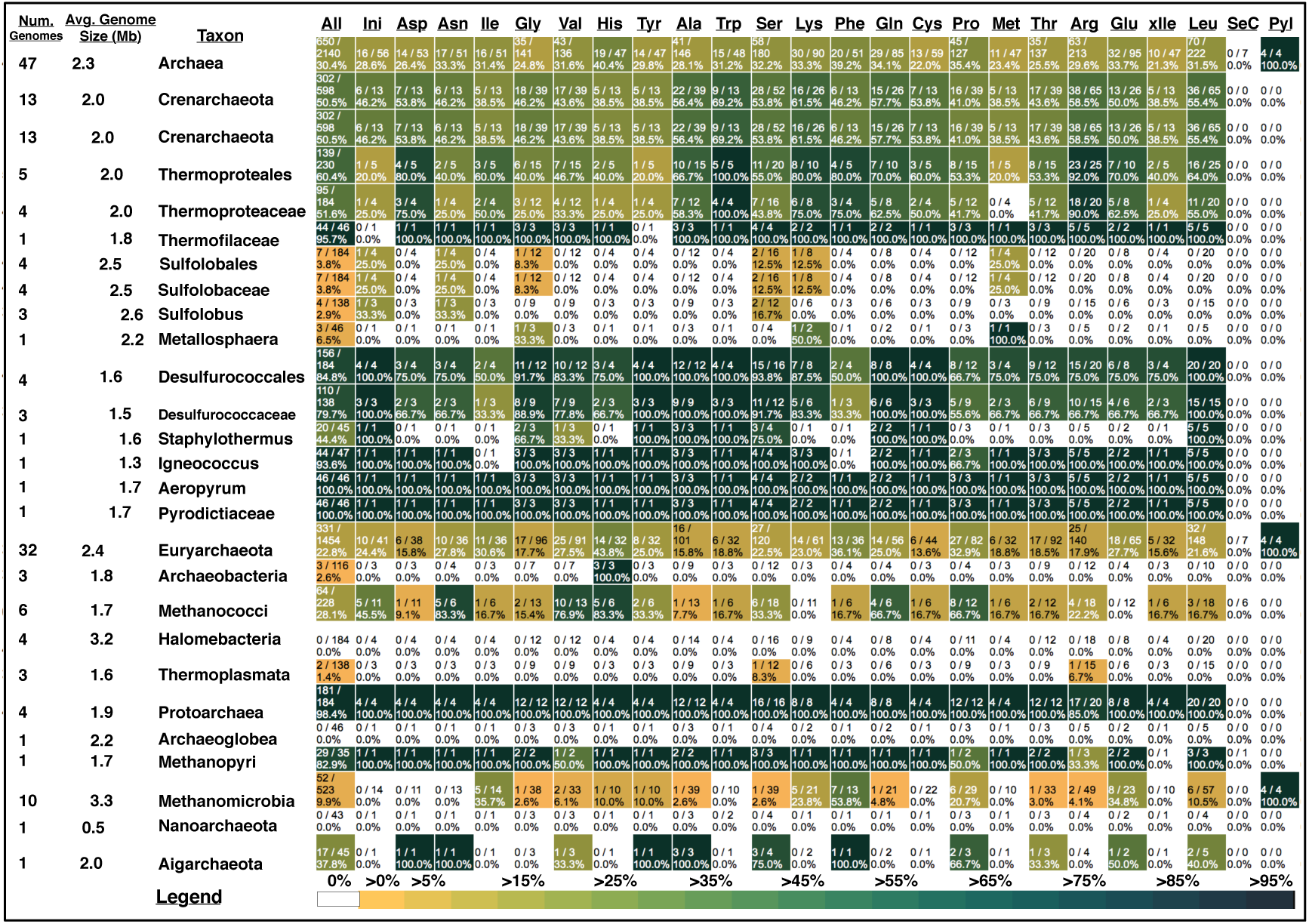
Summarized genome size and CCA frequency data in Archaeal clades broken out by tRNA functional class. Annotations are the same as in Figure 2.

## DISCUSSION

We observed widespread phyletic variation in the frequencies and patterns at which tRNA genes template CCA across functional classes in prokaryotic genomes. Across diverse bacterial and archaeal clades, frequencies range between 0 to 100%. The key finding is that initiator tRNA genes have the greatest class-specific frequency of CCA-templating in bacteria after tRNA^Sec^ genes. Furthermore, in diverse bacterial lineages, especially among the reduced genomes of free-living Cyanobacteria and host-associated endosymbiotic Proteobacteria, initiator tRNA genes template CCA at uniquely high frequencies. In Proteobacteria, all tRNA genes template CCA at high rates, but initiator tRNA genes have the highest overall rate, second only to tRNA^Sec^ genes.

We believe that the tRNA gene reannotations that led to our results are accurate. The most important source of reannotation errors would be from our reclassification of tRNA gene function (Table 1). Note that no previously annotated initiator tRNA genes were reclassified in our analysis, but rather a substantial fraction of genes annotated as elongator tRNA^Met^ were reclassified as tRNA^lMet^ or tRNA^Ile^. Of these reclassifications, detection of initiator tRNA genes by TFAM has very high sensitivity and specificity (Ardell and Andersson 2006, Silva, Belda et al. 2006). This is because initiator tRNA sequence and structure is highly conserved over the three domains of life (Marck and Grosjean 2002).

For example, one of the Cyanobacterial lineages shown in Figure *2* — *Gloeobacter —* is annotated as not having any initiator tRNA genes in tRNAdb-CE v.8. In our analysis of the tRNA gene complements of 2323 prokaryotic genomes in tRNAdb-CE v.8, initiator tRNA genes were not annotated in only 15 genomes (0.6%). We spot-checked several of these aberrant tRNA gene complements by examining their score distributions with TFAM 0.4 to verify that there were no viable candidates for initiator tRNA genes in these gene complements. No tRNA genes in any of the gene complements that we checked scored outside of the normal background distribution for the initiator tFAM model. We believe that initiator tRNA genes may simply be missing from the genome annotations that were aggregated in tRNAdb-CE v.8. Moreover, statistically, our results are robust to these missing data.

The initiator tRNA of protein synthesis in Bacteria is known as tRNA^fMet^, because its charged methionine moiety contains a formyl group attached to the *α*-amino group. By templating CCA in tRNA^fMet^ genes, Bacteria can directly synthesize tRNA^fMet^ with the CCA sequence at the 3’-end. Below we hypothesize five non-mutually exclusive potential advantages for tRNA genes to template CCA.

Our first hypothesis is that certain tRNA classes, particularly initiator tRNAs, may have relatively non-conforming structures that lead to inefficient processing in shared tRNA maturation pathways. For example, tRNA^fMet^ in Bacteria is exceptional in that it contains a mismatched C-A pair at the 1–72 position of the acceptor end, providing a C1-A72 motif for recognition by initiation factors to initiate protein synthesis ((Lee and RajBhandary 1991). All elongator tRNAs contain a Watson-Crick (W-C) base pair at the 1–72 position and therefore are discriminated against by initiation factors. However, the C1-A72 motif of tRNA^fMet^ compromises the efficiency of processing at the 5’-end (Meinnel and Blanquet 1995), so direct transcription of the CCA sequence can be a critical component to mitigate this reduced efficiency (Wegscheid and Hartmann 2006, Wegscheid and Hartmann 2007). In contrast, initiator tRNAs in Archaea have Watson-Crick base-pairs in the 1–72 position (Marck and Grosjean 2002) so we conjecture that they do not share this “Achilles heel” problem with Bacterial initiator tRNAs, but instead are efficiently processed at the 5’-end without any requirement for a 3’-end CCA sequence. This is consistent with our observation that initiator tRNA genes in Archaea do not have a particularly high frequency of CCA-templating.

Second, direct templating of the CCA sequence in tRNAs can potentially increase the maximal growth rate of cells. Under conditions of rapid growth, the co-transcriptional synthesis of 3’-terminal CCA in tRNAs can increase the allocation of cellular resources directly to the synthesis of new proteomic biomass and growth in two ways: first, by reducing or eliminating steady-state cellular pools of species of nonfunctional tRNA precursors, which reduces the mass and energy overhead of the translational machinery itself, and second, by reducing the steady-state fraction of ribosomes devoted to synthesizing tRNA-affiliated proteins such as CCA-adding enzyme (Ehrenberg and Kurland 1984, Klumpp, Scott et al. 2013).

Third, given that translational initiation is rate-limiting in protein synthesis (Vind, Sorensen et al. 1993), and therefore a key determinant of maximal growth rate (Ehrenberg and Kurland 1984, Hersch, Elgamal et al. 2014, Pop, Rouskin et al. 2014), cells selected for a high maximum growth rate may need to efficiently maintain high concentrations of initiator tRNA^fMet^ for rapid growth. The costs of maturation of a tRNA to a growing cell should increase proportionally with the concentration of that tRNA, and initiator tRNA concentration increases more with growth rate in *E. coli* than elongator tRNAs (Dong, Nilsson et al. 1996), so the fitness impact of templating CCA in initiator tRNAs should be greater than in elongator tRNAs in rapidly growing cells. For example, the record-high growth rates reported among *Vibrio* species (Aiyar, Gaal et al. 2002) is associated with very high initiator tRNA gene copy numbers in *Vibrio* genomes (Ardell and Andersson 2006). Consistent with the above, all initiator tRNA genes in *Vibrio* template CCA in the present analysis.

Fourth, rapid synthesis of initiator tRNAs through co-transcriptional synthesis of CCA could reduce the lag phase associated with the transition to growth by reducing the waiting time to increase initiator tRNA concentration. Importantly, this “bootstrapping” trait may be important for all cells, including free-living and endosymbiotic bacteria under reductive genome evolution, and not just for cells capable of rapid growth. Many such cells could have an advantage in the rapid initiation of protein synthesis from a quiescent state in response to environmental change. Indeed, we have shown that each nucleotide addition for post-transcriptional synthesis of CCA requires the CCA enzyme to proofread tRNA integrity (Dupasquier, Kim et al. 2008, Hou 2010), which likely delays maturation of newly transcribed tRNAs.

Fifth, for elongator tRNA genes, direct templating of CCA can facilitate more rapid synthesis of corresponding tRNA elongators to help cells avoid transient depletion of specific ternary complexes and the detrimental consequences that such shortages may have on the accuracy of protein synthesis and proteomic integrity. The supply-demand theory of tRNA charging dynamics (Elf, Nilsson et al. 2003) predicts wide variability in sensitivity of charging levels of tRNA species to perturbations, such as amino acid starvation, affecting specific elongator tRNAs for both proteomically abundant and rare amino acids such as Leucine, Tyrosine and Phenylalanine. Stalled ribosomes caused by shortages of specific ternary complexes increase translational misreading at corresponding “hungry” codons (O’Farrell 1978, Gamper, Masuda et al. 2015), including frame-shift errors (Gallant and Lindsley 1998), all of which can cause protein misfolding, aggregation, and damage (Drummond and Wilke 2009).

While many cyanobacteria with reduced genomes are not fast-growing, they may generally be subject to multiple constraints of chronic nutrient limitation and a heavy burden of a large fraction of proteome dedicated to autotrophic functions (Burnap 2015). When combined, these factors may lead to “proteomic constraints” from small cell sizes, an exacerbation of macromolecular crowding, and increased sensitivity to mistranslation of the most abundant parts of the proteome (Burnap 2015). We suggest that the relatively high frequency at which tRNA^Tyr^ genes template CCA in Cyanobacteria (Fig. 2) is associated with a unique biological sensitivity to depletion of charged tRNA^Tyr^. Tyrosine residues are critically important for both catalysis and stability of RuBisCo (Esquivel, Pinto et al. 2006), one of the most abundant proteins in Cyanobacteria (Wegener, Singh et al. 2010). In light of this hypothesis it is remarkable that there exists a d-Tyr-tRNA^Tyr^ deacylase that is conserved and apparently unique to Cyanobacteria (Wydau, van der Rest et al. 2009), which helps maintain the accuracy of tRNA^Tyr^ charging. Competition experiments that model biologically relevant conditions with Cyanobacterial strains with or without CCA-templating for tRNA^Tyr^, as well as biochemical assays, could test this hypothesis.

We further suggest that the advantages of avoiding supply shortages and streamlining tRNA biogenesis pathways may extend to other elongator tRNAs that we found to template CCA in an often lineage-specific manner. Selenocysteine and Pyrrolysine tRNAs both have complex biosynthetic/maturation pathways and both template CCA at high frequencies in our analysis. Similarly, biosynthesis of Asn-tRNA^Asn^ involves two steps, first by synthesizing a mispaired Asp-tRNA^Asn^, followed by conversion of Asp to Asn (Curnow, Ibba et al. 1996, Becker and Kern 1998, Bailly, Blaise et al. 2007). Indeed, genes for both tRNA^Asp^ and tRNA^Asn^ template CCA at high frequencies.

Although the synthesis of Gln-tRNA^Gln^ also relies on a two-step pathway involving transamidation of Glu on Glu-tRNA^Gln^ (Gagnon, Lacoste et al. 1996), the frequencies for tRNA^Glu^ and tRNA^Gln^ are among the lowest we observed in Bacteria overall. Further analysis and experiments will be necessary to fully understand the patterns reported in this paper.

Re-annotation of tRNA gene sequences was essential to our discovery that CCA-templating is a major feature of initiator tRNA genes. This shows the importance for genome annotation projects of using tRNA gene-finders with taxonomically correct models. More generally, this work demonstrates the importance of using bioinformatic assets carefully to maximize scientific returns.

## MATERIALS AND METHODS

**Data**. Version 8 (October, 2014) of the tRNAdb-CE database (Abe, Inokuchi et al. 2014) was downloaded on November 4, 2014. NCBI Taxonomy data (NCBI Resource Coordinators 2014) was downloaded on November 13, 2014.

**Functional Reannotation of CAU-anticodon tRNAs**. We classified bacterial CAU-anticodon-templating tRNA genes as templating methionine elongators, lysidinylated isoleucine elongators or initiators using TFAM version 1.4 (Ardell and Andersson 2006, Tåquist, Cui et al. 2007) with a general bacterial model for this purpose based on a previously published analysis (Silva, Belda et al. 2006).

**Structural Annotation of 3’*-*ends**. To annotate Sprinzl coordinates to the 3’-end of each tRNAdb-CE sequence record, we implemented a dynamic programming algorithm to optimize base-pairing of the annotated 3’-end of the mature tRNA in each record against its own annotated 5’-end and trailer sequence.

For each sequence record we obtained the 5’-most seven bases of the annotated acceptor stem sequence and reversed it to obtain sequence *x*. Given sequence *x*, we computed its optimal pairing against a second sequence *y* defined by the last 12 bases of the annotated 3’-end and the first five bases of the annotated 3’-trailer using the simple dynamic programming algorithm described here.

Let *x* and *y* be finite sequences over the alphabet Σ = {*A, C, G, U*}, with lengths *m* and *n* respectively. We compute a matrix *H* whose elements are specified as follows:

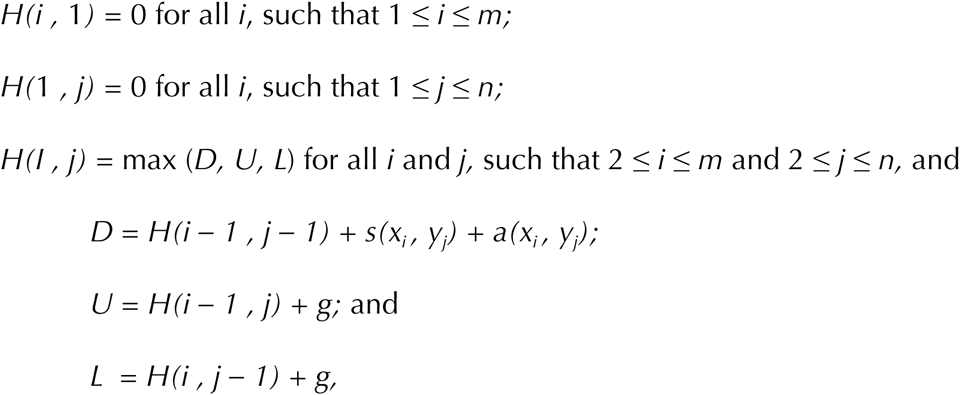

where *x_i_*, is the *i*th base in sequence *x* of length *m = 7, y_j_* is the *j*th base in sequence *y* of length *n = 17, s(x_i_, y_j_) =* 4, for *(x_i_, y_j_) ∈ { (A,U), (U,A), (C,G), (G,C), (G,U), (U,G)}* and *s(x_i_, y_j_) =* 1 otherwise, *a(x_i_, y_j_)* is an annotation bonus if *x_i_*, and *y_j_* were annotated as paired in tRNAdb-CE, *g* = −5 is a linear gap penalty, and *H(i, j)* is the maximum base-pairing score obtained on sequence prefixes *x*[*1,i*] and *y*[*1,j*]. We compared results both with and without an annotation bonus, *i.e*. we recomputed *H(m, n)* for every record using the bonus *a(x_i_, y_j_) = 1* or no bonus *a(x_i_, y_j_) = 0*.

**Statistics and Visualization of Genome Size and CCA-templating Data**. After reannotation, we considered a tRNA gene to template CCA if Sprinzl bases 74 through 76 contained the sequence CCA. We used genome size data downloaded as a “genome report” from NCBI Genome on October 26, 2015 (NCBI Resource Coordinators 2014) and visualized data using the Interactive Tree of Life (Letunic and Bork 2007, Letunic and Bork 2011)

## ACKNOWLEDGEMENTS

We thank the support of NSF grant 1344279 to DHA and NIH grants, 1R01 GM114343, 5U01 GM108972, and 1R01 GM068561 to YMH.

## SUPPLEMENTARY MATERIALS

1. **README.txt** — description of each file.
2. **tCE-Nov5–2014.3.fas** — reannotated tRNAdb-CE v.8 data
3. **prokaryotes_NCBI_102615.txt** — genome size data from NCBI.
4. **addresses_v4.txt.gz** — NCBI taxonomy data required by compute_heatmap.pl
5. **silva.coveam** — TFAM model used to reannotated tRNA functions.
6. **model_cca.pl** — Perl script for structural reannotation of 3’ends.
7. **compute_heatmap.pl** — Perl script to generate full data browser and iTol input.
8. **iTol_color_definitions.txt** — Color definitions for clades to generate Figure 1.
9. **phyloT.tre** — Phylogenetic tree of NCBI taxon IDs from NCBI Taxonomy.
10. **CCA-HEATMAP-TAX-BROWSER/Ardell_Hou.html** — web-browser based navigator of full dataset.
11. **Ardell_Hou_all_data.pdf** — static and searchable PDF with full dataset.

